# Dataset Documentation for Responsible AI: Analysis of Suitability and Usage for Health Datasets

**DOI:** 10.1101/2025.11.18.689064

**Authors:** Anna Heinke, LingLing Huang, Kyongmi U. Simpkins, Fritz Gerald P. Kalaw, Apoorva Karsolia, Kiratjit Singh, Sanjay Soundarajan, Camille Nebeker, Sally L. Baxter, Cecilia S. Lee, Aaron Y. Lee, Bhavesh Patel, the AI-READI Consortium

**Affiliations:** Jacobs Retina Center, 9415 Campus Point Drive, La Jolla, California, United States of America, 92037; Viterbi Family Department of Ophthalmology and Shiley Eye Institute, University of California San Diego, 9415 Campus Point Drive, La Jolla, California, United States of America, 92037; Division of Ophthalmology Informatics and Data Science, Viterbi Family Department of Ophthalmology and Shiley Eye Institute, University of California San Diego, 9415 Campus Point Drive, La Jolla, California, United States of America, 92037; Division of Biomedical Informatics, Department of Medicine, University of California San Diego, La Jolla, California, USA; Department of Electrical and Computer Engineering, University of California Davis, Davis, California, United States of America, 95616; FAIR Data Innovations Hub, California Medical Innovations Institute, San Diego, California, United States of America, 92121; Herbert Wertheim School of Public Health & Human Longevity Science, University of California San Diego La Jolla, California, United States of America, 92093; Department of Ophthalmology, University of Washington, Seattle, Washington, United States of America; John F. Hardesty MD Department of Ophthalmology and Visual Sciences, Washington University in St Louis. Saint Louis, Missouri, United States of America

**Keywords:** Data documentation, Bias, Transparency, Health data, Artificial Intelligence, Machine Learning

## Abstract

Artificial Intelligence (AI) is rapidly transforming healthcare, but also raising concerns about algorithmic biases that mostly stem from the training data. It is widely supported that transparent dataset documentation is key to enabling responsible AI development. Several standardized dataset documentation approaches have been established, such as Datasheet, Dataset Nutrition Label, Accountability Documentation, Healthsheet, and Data Card. However, their suitability and usage for health datasets remain unclear. In this work, we compared all five approaches and evaluated their alignment with the STANDING Together Recommendations for Documentation of Health Datasets. We also investigated their real-world usage and gathered insights from generators and consumers of health datasets. Our findings reveal that none of these documentation approaches are used widely or fully suited for health datasets. We recommend developing a standard documentation approach for health datasets along with clear guidelines and automation tools to support adoption.

---

Artificial Intelligence (AI) is transforming clinical research and healthcare as well as revolutionizing diagnostics and treatment strategies.^1,2^ However, many studies have warned that AI-powered technologies may reinforce existing biases or introduce new ones. A key source of bias in AI models is the data they are trained on.^3–6^ For instance, dermatology AI models used to detect skin conditions often perform worse on dark skin tones and uncommon dermatological diseases because their models were developed using data skewed towards fair skin and common diseases.^7^ Biases can be introduced throughout a dataset’s lifecycle, from study design to data collection and data processing. Contributing factors include underrepresentation of certain groups in the dataset, approaches used to aggregate and label data, and lack of diversity in the study team.^8^

Health datasets used to train AI models often come from clinical care or observational studies and reflect constraints from their original settings, such as the study’s goals, the location of data collection, or the resources available. Therefore, eliminating factors that may lead to algorithmic biases is not always possible for dataset generators. However, they can support the responsible development of AI models by transparently and structurally documenting characteristics such as dataset’s purpose, motivation, recommended use, and attributes, thereby enabling downstream users to better assess potential limitations and sources of bias.^9,10^ Several dataset documentation approaches have been proposed to promote transparency, such as Datasheet^9^, Dataset Nutrition Label^11,12^, Accountability Documentation^13^, Healthsheet^14^, and Data Card.^15^ Their goals are twofold: to encourage dataset generators to reflect carefully on the process of creating a dataset, and to equip dataset consumers with the information needed to make informed use of a dataset for their AI-related tasks.

This work aims to explore the suitability and usage of these approaches for documenting health datasets. Here, a health dataset refers to any set of data collected from human subjects for a health-related study. To our knowledge, there are currently no community-accepted guidelines for the documentation approach to use for health datasets. The STANDING (STANdards for data Diversity, INclusivity and Generalisability) Together program has established a set of 18 recommendations for documenting health datasets to address concerns about bias, underrepresentation, and inequity in health data used for AI development.^10,16^ This program represents the most comprehensive effort for guiding data generators. However, they do not provide guidance on the structure of dataset documentation. Rather, they “invite those using these recommendations to consider existing artifacts when structuring their documentation, including Datasheets for Datasets or Healthsheets.” Without clear guidelines, it could be difficult for data generators to select the right documentation approach. We experienced this challenge firsthand while deciding on a standardized documentation approach for the AI-READI (Artificial Intelligence Ready and Exploratory Atlas for Diabetes Insights) dataset, a multimodal dataset being created by the National Institutes of Health (NIH) Bridge2AI Program.^17^ We eventually decided to create a Healthsheet due to its health-oriented focus, though without an extended understanding of the different documentation approaches and their usage in the field.

Here, we reviewed the different documentation approaches to understand how they align with the needs of health datasets. Specifically, we examined their motivations and content to identify areas of overlap and differences. We also analyzed how they align with STANDING Together Recommendations for Documentation of Health Datasets.^10^ We then investigated the real-world adoption of these approaches and analyzed their usage for health datasets. Finally, we conducted a survey among generators and consumers of health datasets regarding their awareness, usage, and perceptions of these documentation approaches. Our goal was to provide insights that inform best practices for documenting health datasets.

## Results

### Review of the documentation approaches

Based on our knowledge and discussions with colleagues, we identified five main approaches to prepare human-friendly dataset documentation for responsible AI: 1) Datasheet, 2) Dataset Nutrition Label, 3) Accountability Documentation, 4) Healthsheet, and 5) Data Card. We reviewed associated papers and resources to understand when they were established, their goals, and the methods for establishing them.^9,11–15,18,19^ Their key characteristics are summarized in **Table 1**, and an overview is provided below in the chronological order of their development.

**Table 1.**
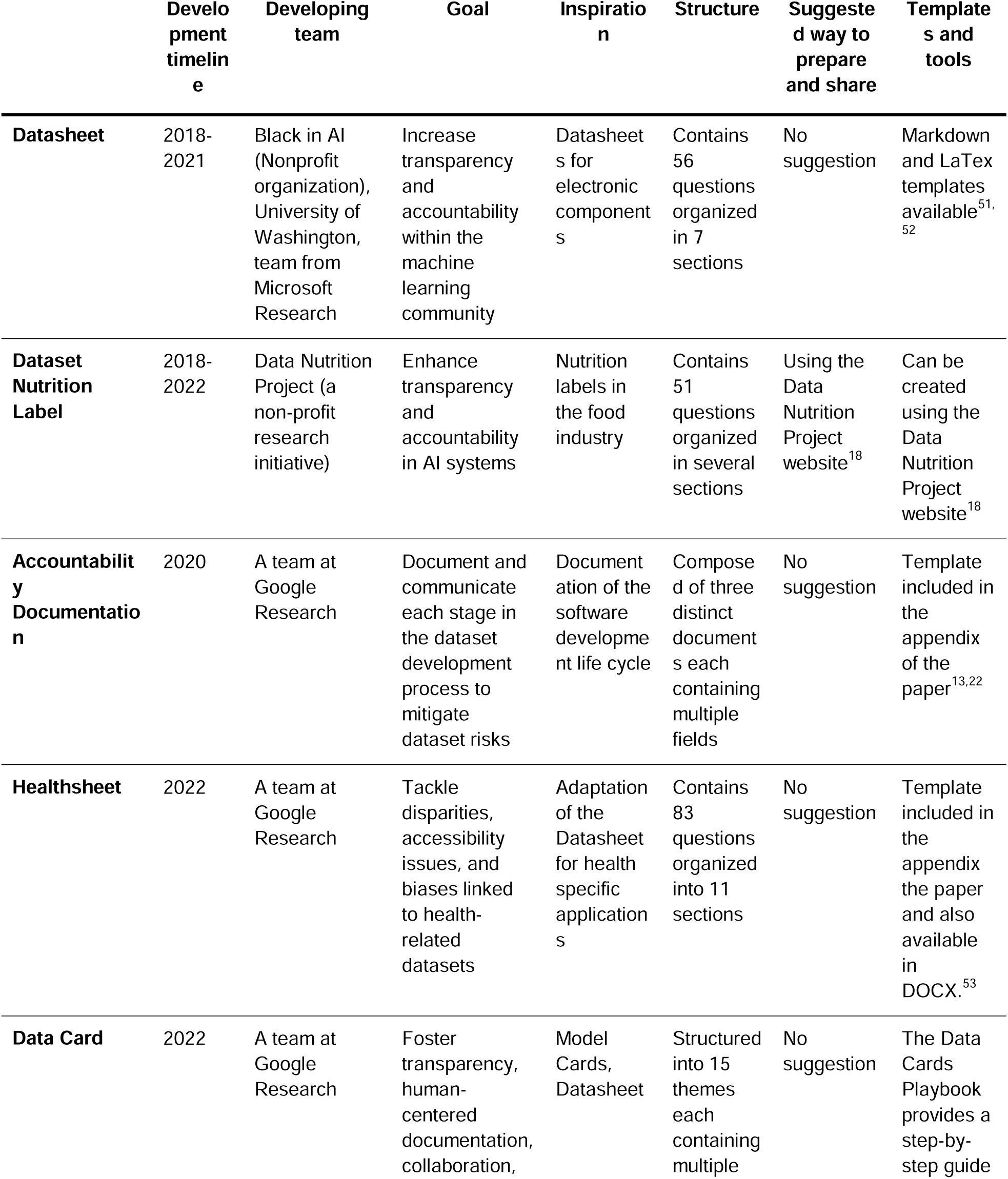
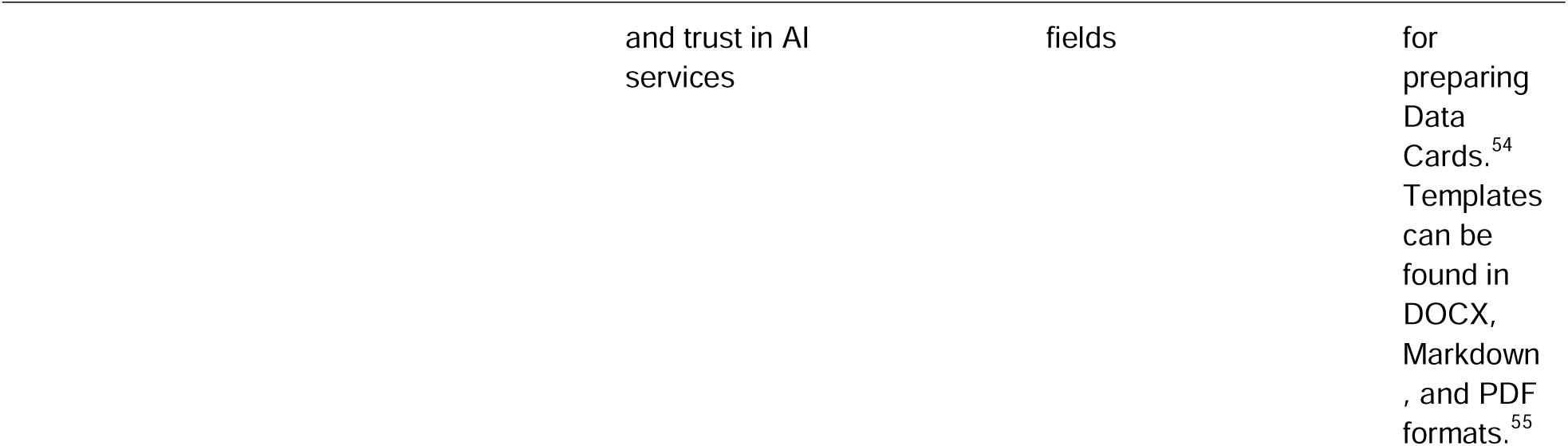
Overview of the different dataset documentation approaches analyzed in this work.

The Datasheet for datasets was inspired by Datasheets used in the electronics industry for documenting electronic components. It is designed as a series of questions. Its first draft was introduced in a preprint in 2018.^20^ Several versions were then shared through preprints before a final version was eventually published in December 2021.^9^ The Dataset Nutrition Label was inspired by nutrition labels used for food and beverages. Just as food nutrition labels summarize essential information about what you consume, the Dataset Nutrition Label was designed to summarize essential facts about a dataset that will be used for AI/ML developments. The first generation Dataset Nutrition Label was introduced in 2018, followed by the second generation in 2020 and the third generation in 2022.^11,12,21^ The first generation introduced a blend of qualitative and quantitative modules supported by various statistical and probabilistic models. The second and third generations shifted towards a more user-centered approach, providing key information about a dataset through interactive panes. The label presents a visually appealing snapshot of the dataset, enhanced by icons and symbols. Another approach is what we will refer to as Accountability Documentation, as there is no official name. This approach is inspired by software development lifecycle practices. It is designed as a set of three documents each containing multiple fields to document information about each stage in the dataset development process. An initial preprint about the framework was first shared in October 2020 and later published in March 2021.^13,22^ The Healthsheet is an adaptation of the Datasheet for health-specific applications. It was introduced in a preprint in February 2022 and was later published in June 2022.^14,23^ The Data Card is a structured summary of essential information about a dataset. It was introduced in a preprint in April 2022 that was later published in June 2022.^15,24^ According to the authors, the Data Card is aimed at broader understandability for diverse readers, promoting informed decision making about data usage for products, research, and policy. It is designed with a flexible, modular structure that can be extended as needed for different datasets.

Overall, these five documentation approaches share the common goal of enhancing transparency and accountability in AI development by providing structured information about datasets. Each approach aims to support better decision making by dataset generators and dataset consumers. Some have even inspired each other. The Healthsheet is a direct adaption of the Datasheet for health datasets, and the authors of the Dataset Nutrition Label state that they “have pulled many questions directly from the thorough and thoughtful work of the Datasheets for Datasets project”.^12^ However, all five approaches still differ from each other in focus and structure. The Datasheet and Healthsheet encourage critical reflection through detailed, often open-ended questions. The Dataset Nutrition Label takes a more visual and interactive approach, designed to be user-friendly and to appeal to all audiences. The Accountability Documentation uniquely applies a lifecycle model, while the Data Card offers a concise, modular structure for describing datasets at a very granular level. Distinctively, only the Healthsheet was specifically developed for health datasets. Its authors note that existing documentation approaches like Datasheets do not fully address the complexities and ethical considerations specific to health data, which motivated the development of the Healthsheet as a more suitable alternative.^25^ The Dataset Nutrition Label is expected to be prepared and shared using its dedicated platform. For all other documentation approaches, neither the format (e.g., PDF or Markdown) nor the method of sharing (with the dataset or separately) is specified.

### Comparison of the content

We compared the information documented by the different dataset documentation approaches to understand their similarities and differences. To achieve that, we used their respective templates.^9,13,14,18,19,26^ Direct comparison was challenging because different approaches often asked for similar information using different wording or structure. To enable systematic comparison, we manually curated each question/field from a data documentation approach into what we called metadata elements, i.e. simple sentences describing the information requested. When a question or field asked for multiple distinct pieces of information, we split it into several metadata elements. Conversely, when different questions asked for the same high-level information, we assigned the same metadata element to each of them. We made sure to use the same metadata element for similar information across the different data documentation. This process is illustrated in **Fig. 1**. We similarly broke down the STANDING Together recommendations^10^ into metadata elements to analyze how the content of the dataset documentation approaches aligns with these recommendations.

**Figure 1.**
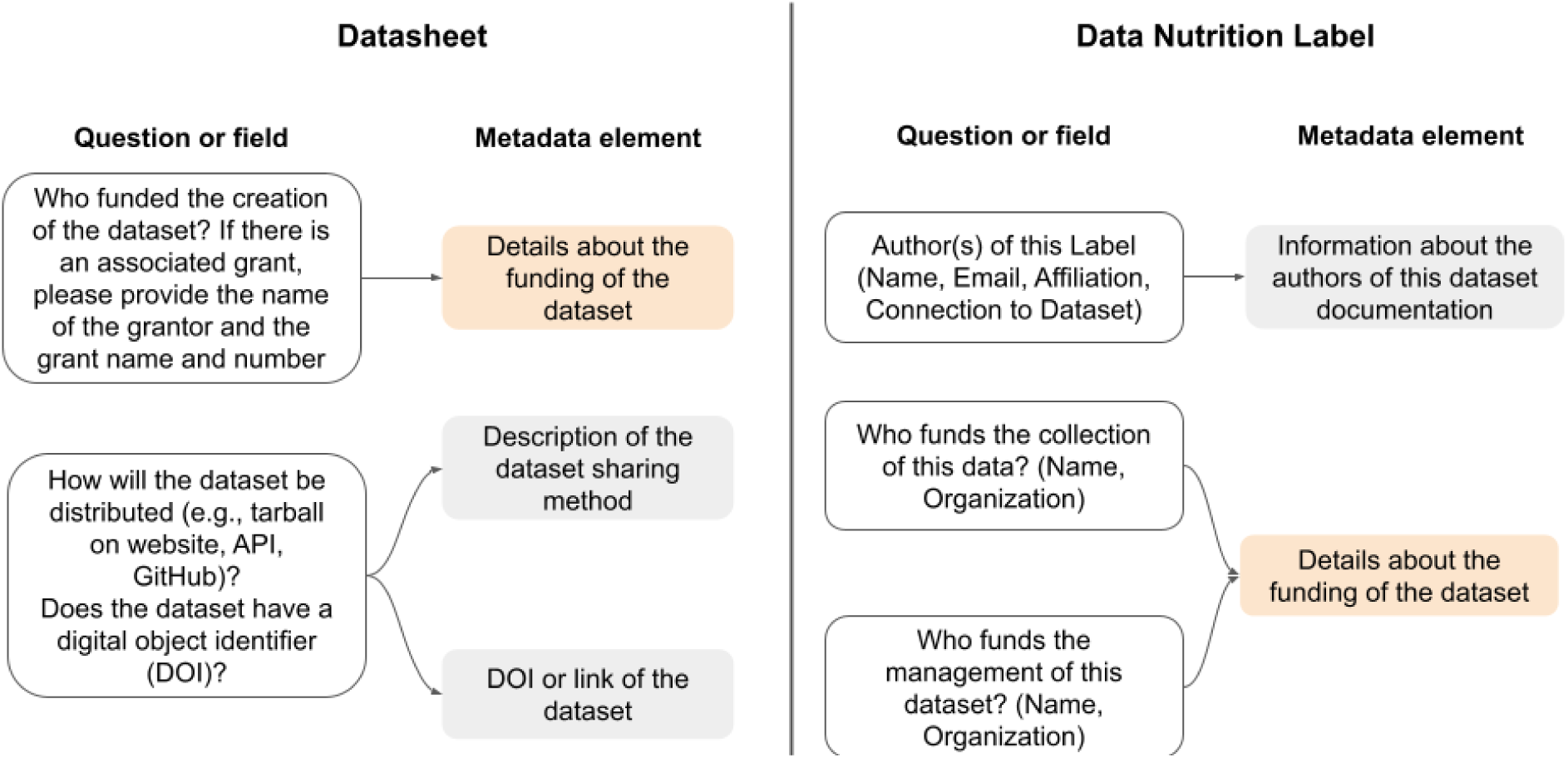
Illustration of the approach to assign metadata elements to each question field of the different dataset documentation approach. We took each question/field from a data documentation approach and described what information it is asking in a simple sentence that we called metadata element. When multiple distinct information was asked in a given question/field we split it into multiple metadata elements. In other cases, we also assigned the same metadata element to multiple questions/fields when they were asking the same high-level information. We made sure to use the same metadata element for similar information across the different data documentation (for example, “Details about the funding of the dataset” in this illustration).

We identified 138 unique metadata elements across all five data documentation approaches (**Fig. 2**), representing 138 unique pieces of information documented across them. A full list of the metadata elements identified is provided in the dataset associated with this work (see **Data Availability** section). Of them, 76 (or 55%) are found exclusively in one of the five documentation approaches. The Datasheet has only one exclusive metadata element out of 47 (2%) related to documenting the risk of identifying individuals in the dataset. The Dataset Nutrition Label has 14 exclusive metadata elements out of 43 (33%). Specifically, it is the only approach that requires documenting details about the creation of the dataset documentation, including information about the creators of the data documentation, their familiarity with the dataset, and who was consulted for creating it. Other exclusive metadata elements include keywords, information about the format/structure of the dataset, and the required domain-specific knowledge for proper use of the dataset. The Accountability Documentation has 21 exclusive metadata elements out of 49 (43%). It is the only approach that focuses on documenting the ethical impact of the data and its collection, data collection cost, and details about related datasets and documents. The Healthsheet has 12 exclusive metadata elements out of 67 (18%) that mostly cover information relevant to health-related datasets, such as the number of subjects represented in the dataset, study inclusion criteria, language(s) used to communicate with study participants, countries where data were collected, and description of the strategies to avoid reidentification. Finally, the Data Card has 28 exclusive metadata elements out of 53 (53%), including detailed statistics for each data field, handling of outliers, dataset validation processes, dataset annotation, and alignment with upstream sources.

**Figure 2.**
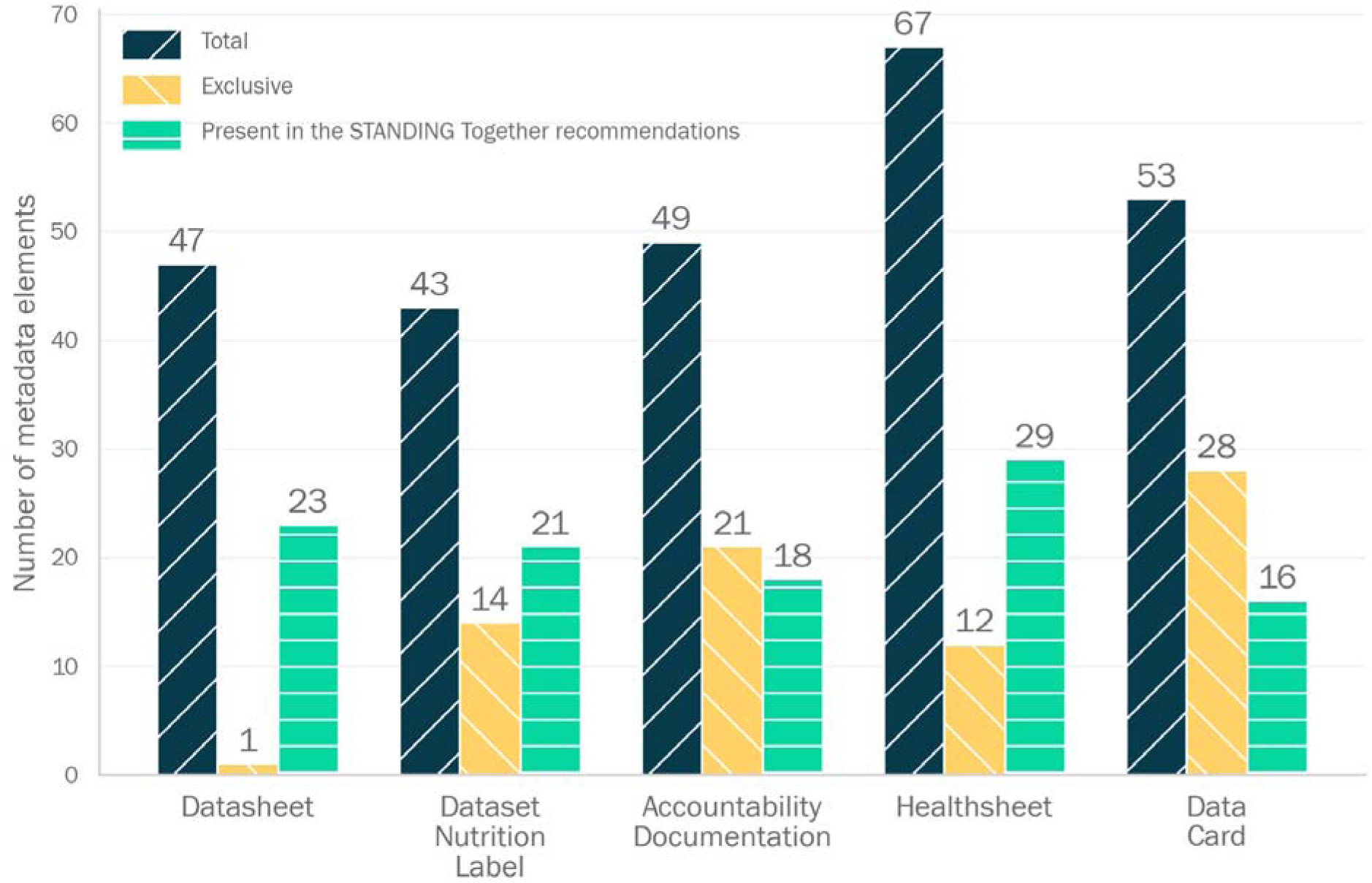
Number of metadata elements identified in each dataset documentation approach. For each documentation approach three numbers are shown: 1) The number of total metadata elements, 2) The number of metadata elements that are exclusive to a dataset documentation approach meaning they are not found in any other, and 3) The number of metadata elements that are similar to one of the 51 identified in the STANDING Together’s Recommendations for Documentation of Health Datasets.

When comparing the metadata elements across the documentation approaches (**Fig. 3**), we find that the Datasheet and Healthsheet are the most similar, with 98% of the metadata elements in the Datasheet found in the Healthsheet and 69% of the metadata elements in the Healthsheet found in the Datasheet. This reflects how the Healthsheet was developed by taking the Datasheet and adding new elements. The Datasheet and Dataset Nutrition Label also have noticeable overlap, given that questions from the Datasheet were included in the Dataset Nutrition Label starting from the second generation. Consequently, the Dataset Nutrition Label and Healthsheet also have noticeable overlap. All other pairs have less than 50% overlap.

**Figure 3.**
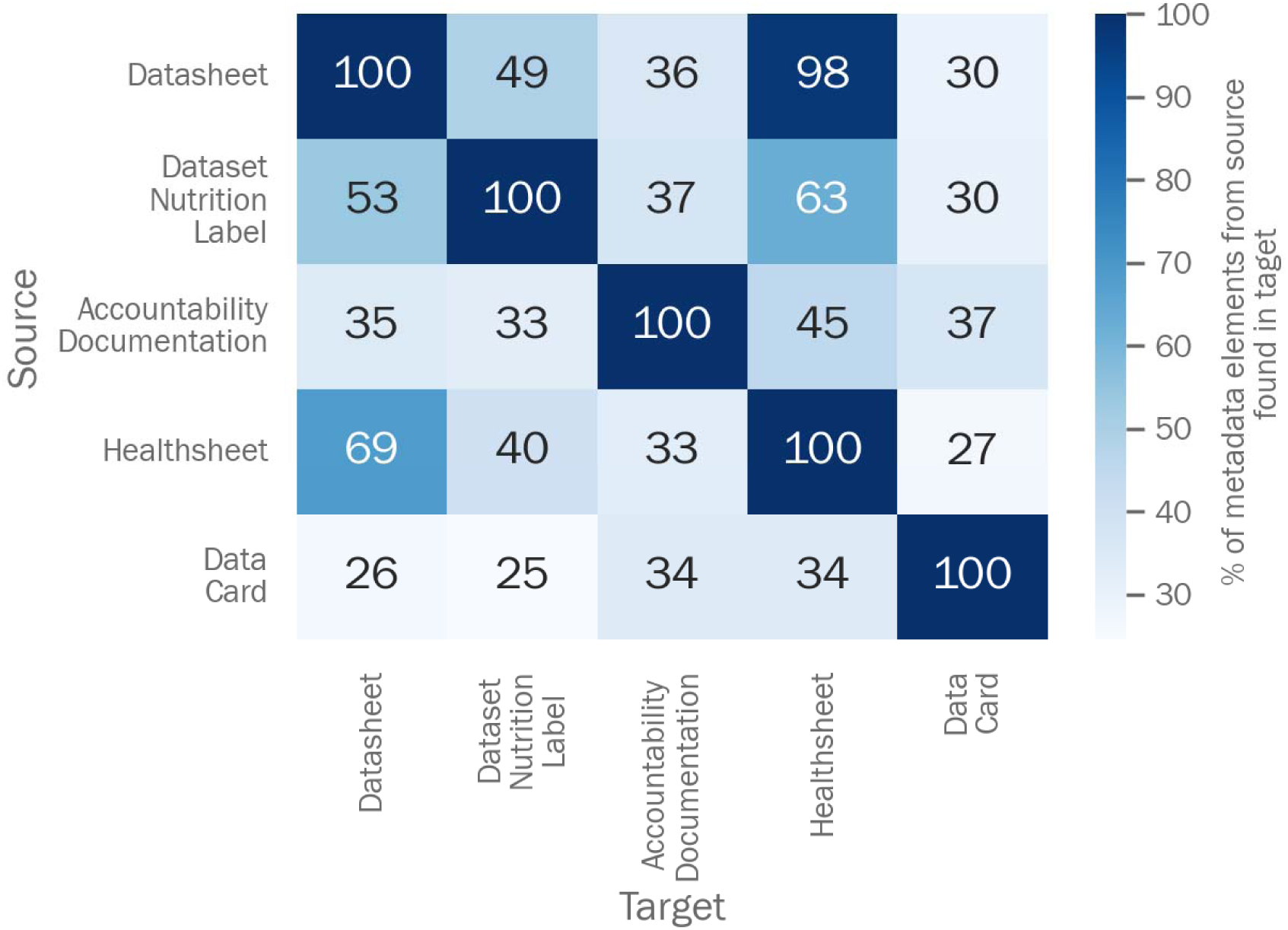
Percent of metadata elements from the source dataset documentation approach found in the target dataset documentation approach.

We identified 51 unique metadata elements in the STANDING Together recommendation. The Healthsheet includes 29 of the 51 metadata elements (57%), which is the most, while the Data Card includes the least with 16 (31%). Thirteen of the 51 metadata elements are absent from all five documentation approaches, covering information such as competing interests amongst dataset creators, description of attempts to mitigate bias, and description of potential bias introduced by the data labelling process.

These results demonstrate that, while there is some overlap, each documentation approach is designed to capture different aspects of a dataset, with certain details exclusive and specialized to each type. However, no single approach fully encompasses all of the recommendations outlined by the STANDING Together initiative, and some recommendations are not addressed by any of these frameworks.

### Usage and adoption of the different approaches

To evaluate how widely these dataset documentation approaches have been adopted, we searched their real-world use in documenting actual datasets. Our strategy consisted of the following four methods: 1) Reviewing the main papers and resources associated with each dataset documentation approach, 2) Reviewing the papers citing their main papers, 3) Searching on GitHub, and 4) Conducting other non-targeted searches. We identified 3,077 resources through these methods and manually screened 919 of them after excluding non-relevant resources through an automated keyword-based screening.

In total, we found 260 dataset documentations that document 254 datasets (some datasets have multiple documentations). The full list is included in the dataset associated with this work (see “**Data Availability**” section). As showcased in **Fig. 4A**, we found 176 Datasheets, 55 Data Cards, 24 Dataset Nutrition Labels, and 2 Healthsheets, while we did not find any use of the Accountability Documentation. We found that the Datasheet has been used to describe 11 health datasets, the Dataset Nutrition Label for 4, the Healthsheet for 2, and the Data Card for 1.

**Figure 4.**
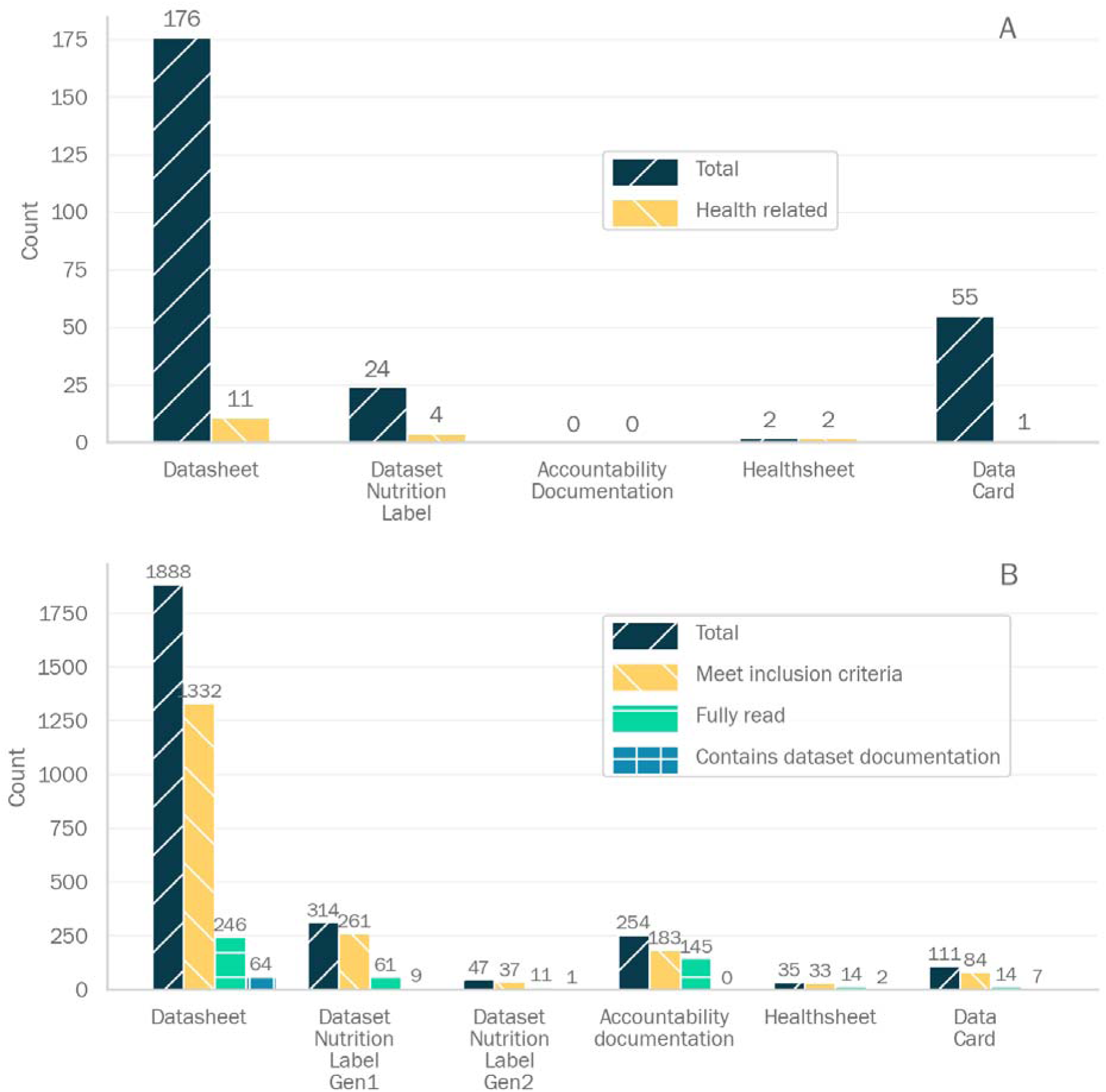
A: Number of real-world usage of the different dataset documentation approaches, including how many were created to document health-related datasets. B: Number of citations to each paper of the different dataset documentation approaches, how many met our inclusion criteria for review, how many were fully read after keyword-based exclusion, and how many of the fully read contained or referred to the dataset documentation approach it was citing.

Through search method 2, we found that only a fraction of the citations to the main paper of each data documentation approach are resources that use the documentation approach for documenting a dataset. For instance, of the 1888 resources citing the Datasheet paper, we were able to access and analyze 1332 because they met our inclusion criteria. We manually screened 246 of them because they included Datasheet-related keywords twice or more, and only 64 of them contained or mentioned a Datasheet, which is less than 5% of all the analyzed resources and less than 26% of the manually screened ones (**Fig. 4B**).

Overall, these findings indicate that despite their high citation counts and a recognized need for supporting responsible AI development, use of these dataset documentation approaches remains low and is nearly absent for health datasets.

### Survey finding

To further understand the usage and usefulness of these data documentation approaches in health datasets, we surveyed two categories of domain experts: 1) Data generators, i.e., individuals who collect or generate health data, and 2) Data consumers, i.e., individuals who use health datasets for developing AI models. The survey was designed to evaluate three major topics: 1) Familiarity with each dataset documentation approach, 2) Perceived difficulty of preparing a dataset documentation by data generators or the usefulness of a dataset documentation by data consumers, and 3) Information necessary for responsibly developing AI models that is missing from each dataset documentation approach. Participants were first asked some general questions about their background and familiarity with the different dataset documentation approaches before being randomly assigned one of them for a more in-depth evaluation. Data generators were asked to respond to questions related to difficulty in preparing their assigned documentation approach, data consumers were asked to evaluate questions related to usefulness, and those who identified as both were asked to provide both perspectives. The evaluation was conducted not only on the documentation approach as a whole but also on its specific sections. Due to the limited number of participants, statistical significance was not evaluated. A copy of the survey, the raw responses of the participants, a table summarizing the profile of the survey participants, and tables summarizing the major results are provided in the dataset associated with this work (see **Data Availability** section).

Twenty-nine participants completed the survey: 5 (17%) identifying as data generators, 14 (48%) as data consumers, and 10 (35%) as both. Most participants (86%) had previously heard about at least one type of dataset documentation approach (see **Fig. 5**). However, only 62% had experience with preparing or using any type of dataset documentation. The Datasheet was the most well-known documentation approach amongst the participants, with 27 (90%) responding that they have heard of it and 17 (59%) prepared or used it. In contrast, the Accountability Documentation was the least popular, as only 6 (21%) participants had heard of it and 3 (10%) prepared or used it.

**Figure 5.**
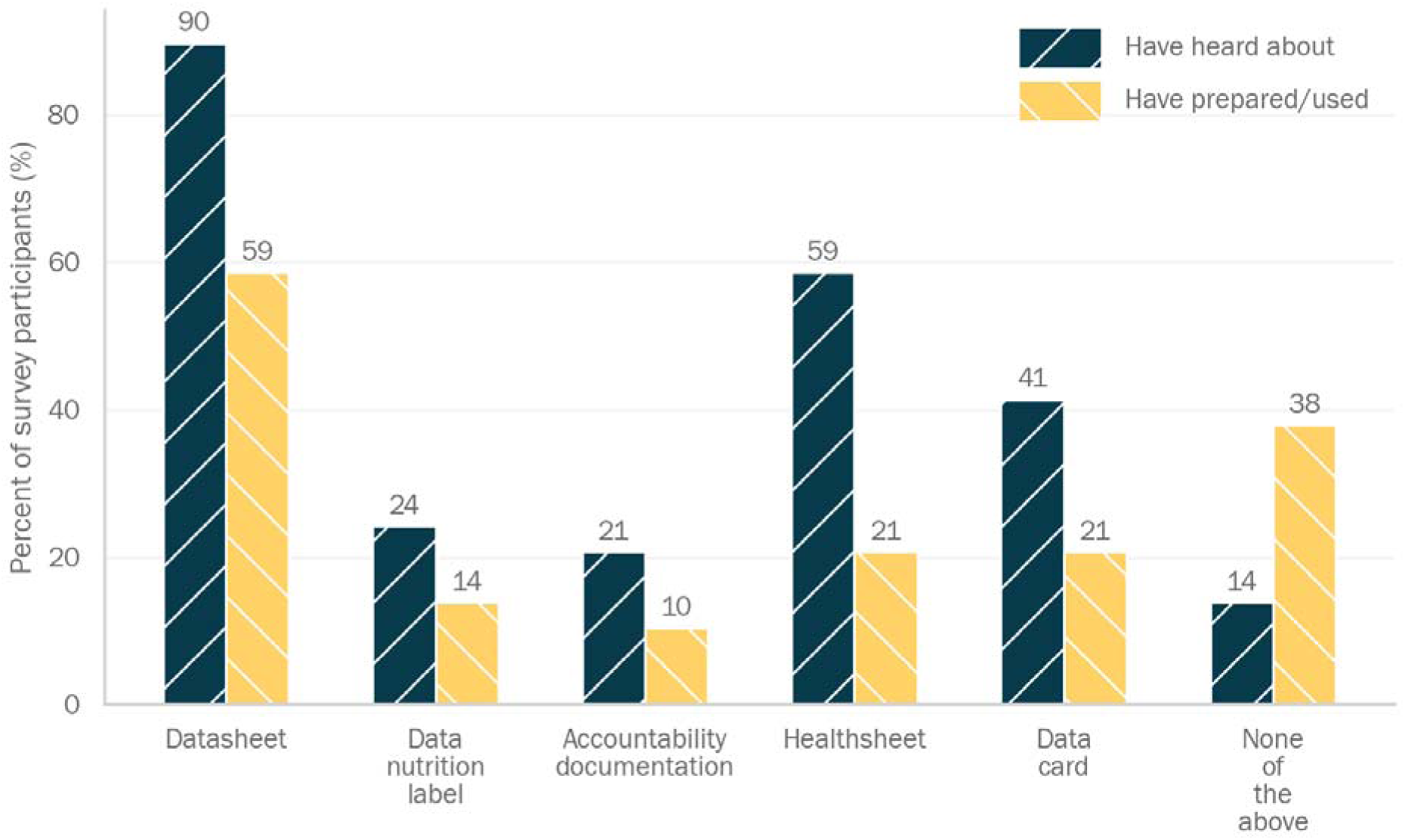
Response from the survey participants when asked “Have you heard about any of these data documentation approaches?” and “Have you used or prepared any of these data documentation approaches?”.

A total of 24 unique evaluations (between 4 and 6 for each documentation approach) of the ease of preparing the documentation approaches were completed by the 5 participants who identified as data generators and the 10 who identified as both. Most evaluations (10/24, 42%) felt “Neutral” regarding the overall difficulty of preparing a given data documentation, followed by “Difficult” (6, 25%). The Datasheet received the highest proportions (3/5, 60%) of “Easy” or “Very easy” to prepare. All other documentation approaches had less than 40% of their evaluations rating them as “Easy” or “Very easy” to prepare, while none of the 4 Accountability Documentation evaluations rated it as such. For section-specific evaluations, we found that all the dataset documentation approaches had multiple sections that were rated “Difficult” or “Very Difficult” by at least one evaluation. Notably, the Accountability Documentation, Data Nutrition Label, and Data Card had one or more sections rated “Difficult” or “Very Difficult” to prepare by 50% or more of their evaluations.

Amongst the 14 data consumers and the 10 participants who identified as both, 45 unique evaluations (between 7 and 11 for each documentation approach) were completed on the usefulness of the documentation approaches in responsibly developing AI models. Most evaluations (37/45, 82%) rated the overall usefulness of a given data documentation as “Useful” or “Very useful”. All documentation approaches had >70% of the evaluations rating them as “Useful” or “Very useful”, with the Healthsheet having 100% and the Datasheet 90%. For section-specific ratings, all dataset documentation approaches had a majority of their sections rated “Useful” or “Very useful” by 50% or more of their evaluations. Notably, all sections of the Datasheet and the Data Nutrition Label met that threshold.

When asked what is missing to enable the responsible development of AI, participants highlighted several gaps in the documentation approaches they evaluated. For the Datasheet, participants noted missing details on comprehensive demographics, dataset versioning, sampling processes, data bias and fairness considerations, model-specific usage warnings, dataset usage history, and mechanisms for user feedback or issue reporting, as well as documentation of IRB protocols and consent procedures. The Dataset Nutrition Label was seen as lacking information on bias identification and mitigation strategies, detailed demographic breakdowns, ethical approvals, informed consent, environmental and social context, reasons for missing data, and clear presentation of data distribution metrics. For the Accountability Documentation, participants pointed out the absence of contextual references, core transparency elements like demographics and instrumentation, explicit bias mitigation measures, and ethical compliance testing. The Healthsheet was found to lack thorough reporting on missing data patterns, bias mitigation steps, ethical approvals, consent processes, data lineage, security considerations, and clear guidance on appropriate and inappropriate dataset use. Finally, participants felt the Data Card did not sufficiently cover social determinants of health, protections for vulnerable populations, dataset generalizability and limitations, ethical review and consent details, sources of potential bias, citation guidance, and documentation of missing data and reasons.

Overall, dataset consumers generally found these dataset documentation approaches useful for responsibly developing AI models, while dataset generators found some of their sections difficult to prepare. Both identified gaps in the information documented in the different dataset documentation approaches.

## Discussion

In this work, we provide a comprehensive evaluation of the Datasheet, Data Nutrition Label, Accountability Documentation, Healthsheet, and Data Card with a focus on their relevance for health datasets. Across our review, comparison, real-world usage analysis, and survey responses, we identified several important patterns and gaps.

Our review and comparison of their content revealed that all five documentation approaches share the common goal of enhancing transparency and accountability in AI development, but differ in their structure, focus, and alignment with health-specific requirements. The Datasheet is designed with open-ended questions that result in an interview-style documentation. Inspired by the Datasheet, the Healthsheet takes the same form but with additional questions tailored for health datasets. The Dataset Nutrition Label is designed to provide a more visual, easy-to-consume overview of a dataset. Drawing from software engineering practices, the Accountability Documentation emphasizes the documentation of a dataset throughout its lifecycle. The Data Card provides a modular and structured summary of a dataset. Our comparison of the content of these documentation approaches provided a more precise picture of their similarities and differences. All of them have overlapping content, but they each also cover unique aspects. The Datasheet contains only one unique metadata element related to the risk of reidentification. The Dataset Nutrition Label stands out for its emphasis on transparency about the dataset documentation process itself. The Accountability Documentation offers a distinct focus on transparency, uniquely documenting information such as the cost of data collection. The Healthsheet integrates health-specific elements, including the number of subjects and study inclusion criteria. The Data Card contains unique details around dataset-level statistics and other technical information relevant for AI model development. Overall, this shows that each documentation approach brings unique aspects that are suited for particular domains or use cases.

The STANDING Together’s effort to provide guidance for transparent documentation of health datasets is, to our knowledge, the largest of its kind to tackle biases in AI health technologies. We found that none of the documentation approaches fully comply with these recommendations. The Healthsheet, though designed for health datasets, covers less than 60% of the recommended information. Even jointly, these documentation approaches fail to address key recommendations such as competing interests, explicit descriptions of bias mitigation efforts, and detailed assessments of potential bias introduced during data labeling. This highlights the need for either a new health-specific documentation approach or guidance for tailoring and synchronizing existing ones to meet these critical recommendations.

Although the documentation approaches studied here are highly cited, our analysis shows their real-world application remains limited and is close to non-existent for health datasets. The Datasheet has seen the broadest adoption, yet only a small fraction of citing papers use one in practice. This discrepancy between recognition and practical uptake for health datasets could be due to the lack of explicit guidelines and support for preparing these dataset documentation.

Our survey of generators and consumers of health datasets reinforces these findings. Participants across roles valued the documentation approaches, recognizing their potential to support responsible AI development. However, their preparation was not universally perceived as easy by dataset generators. Based on our personal experience with documenting the AI-READI dataset, preparing such documentation can be labor-intensive, and there are not currently strong incentives to engage in these efforts. These insights suggest that these documentation approaches, while helpful for responsible AI development, need support to be easy to prepare for dataset generators to support adoption.

While our analysis provides a detailed evaluation of existing dataset documentation approaches, several limitations need to be acknowledged. The process of breaking down the content of the documentation approaches and the STANDING Together recommendations into metadata elements involves interpretative judgement and may not capture all nuances in how the documentation approaches are presented and intended. Moreover, our real-world usage analysis focused on published citations and GitHub, which may underrepresent uses of these documentation approaches, e.g., in industry or internal academic settings. Finally, our survey sample is small and may not capture the variety of perspectives across generators and consumers of health data.

It is widely recognized that thorough documentation of datasets is critical to mitigate biases in AI models stemming from their training data. Together, the findings of this work highlight several key directions for the biomedical field to ensure that health datasets are consistently and effectively documented to support the responsible development of AI models. First, there is a pressing need to establish a standard documentation approach that fully covers the requirements of health data, in particular those conveyed in the STANDING Together recommendations. Second, broader adoption of dataset documentation practices will require increased awareness and incentives. Clear guidelines by funders, data repositories, and scientific journals will be needed to achieve that. Finally, tools that reduce the burden and complexity associated with the preparation of dataset documentation will be essential to make it a common practice in the health AI ecosystem.

## Data availability

The data associated with this manuscript consists of several Excel files (mentioned in the Methods and Results section). Since no FAIR guidelines were found for structuring such data, we structured it according to the SPARC Data Structure (SDS), which provides a broad data and metadata structure to organize biomedical research data in line with the FAIR principles.^27^ The SPARC data curation software SODA for SPARC was used to organize the data and prepare the metadata files.^28,29^ The dataset is maintained in a GitHub repository called “dataset-documentation-paper-data” in the AI-READI GitHub organization, and the version associated with this manuscript (v1.0.0) is also archived on Zenodo.^30^ This data is shared under the permissible Creative Commons Attribution 4.0 International (CC-BY) license.

## Code availability

The code associated with this manuscript consists of several Jupyter notebooks (mentioned in the Methods section). These notebooks are maintained in a GitHub repository called “dataset-documentation-paper-code” in the AI-READI GitHub organization. The notebooks were made FAIR according to the FAIR-BioRS guidelines using Codefair.^31,32^ Accordingly, the version of the code associated with this manuscript (v1.0.0) is archived on Zenodo.^33^ The notebook and associated files are shared under the permissible MIT license.

## Acknowledgement

This work was supported by the US National Institutes of Health (NIH) through grant OT2OD032644.

## Methods

### Review of dataset documentation approaches for responsible AI

Based on our knowledge and discussions with colleagues, we identified five main approaches to prepare human-friendly dataset documentation for responsible AI: Datasheet, Dataset Nutrition Label, Accountability Documentation, Healthsheet, and Data Card. We reviewed associated papers and resources to understand when they were established, their goals, and the methods for establishing them. For Datasheet, Healthsheet, and Accountability Documentation, we reviewed their respective published papers.^9,13,14^ We reviewed the preprints on the first and second generation as well as the Data Nutrition Project website for the Dataset Nutrition Label.^11,12,18^ We reviewed the published paper and the Data Cards Playbook website for Data Card.^15,19^

The HuggingFace dataset card is another form of documentation, used mostly on HuggingFace but we did not consider it in this study because it is intended to be more of a quick summary than a standardized documentation to enable responsible AI development.^34^ We also did not consider domain-specific documentation approaches such as Artsheet, which were not relevant for health datasets.^35^ Note that the focus in this work was on human-friendly documentation for responsible AI development, rather than machine-friendly format.^36–38^

### Comparison of the different documentation approaches

Comparing the information documented by the different dataset documentation approaches is necessary to understand if they are similar or differ greatly to a point where the choice of the documentation approach needs to consider what is deemed important to document for a given dataset. To achieve that, we used their respective templates. For Datasheet, Healthsheet, and Accountability Documentation, we used the templates provided in their respective papers.^9,13,14^ For the Dataset Nutrition Label, we used the Google Doc template from the Data Nutrition Project website.^18^ For Data Card, we used the template provided in the Data Card Playbook and complemented it with additional instructions available in the extended template provided by the Data Card team on GitHub.^19,26^ A direct comparison was found to be difficult because we observed that some of these documentation approaches asked for the same information but in different ways. To enable comparison, we took each question/field from a data documentation approach and described what information it is asking in a simple sentence that we called a metadata element. When multiple distinct pieces of information were asked in a given question or field, we split it into multiple metadata elements. In other cases, we also assigned the same metadata element to multiple questions/fields when they were asking the same high-level information. We made sure to use the same metadata element for similar information across the different data documentation. This process for assigning metadata elements is illustrated in **Fig. 1**. We similarly broke down the STANDING Together recommendations^10^ into metadata elements to analyze how the content of the dataset documentation approaches aligns with these critical recommendations. To verify that each metadata element was unique, we developed a Python-based script in the form of a Jupyter notebook called comparison.ipynb (see the **Code Availability** section) to create a list of all unique metadata elements. We then used the all-MiniLM-L6-v2 sentence transformer (which is based on the MiniLM-L12-H384-uncased model) for embedding them and computed their cosine similarity value.^39–42^ We manually reviewed each pair of descriptions in descending order of cosine similarity value. We noticed that below a similarity score of 0.75, the descriptions were not similar enough, and therefore we did not manually check below that. We identified and merged the metadata elements that were describing the same content but expressed differently, either because of the usage of a different wording or the presence of a typo. are provided in the metadata-elements.xlsx file included in the dataset associated with this work (see the **Data Availability** section).

We then compared these metadata elements across the data documentation approaches to evaluate the similarities and differences between them. This was done with Python code added to the same comparison.ipynb Jupyter notebook. Packages such as pandas^43,44^, Matplotlib^45^, seaborn^46^, and Numpy^47^ were used for all the analysis. One element of our comparison to consider is the customizable nature of some of these data documentation approaches, where it is typically up to the authors to provide the information they deem necessary. In particular, the Data Card is very customizable, so it is, in theory, possible that a Data Card contains all the information asked for in another data documentation approach. For the purpose of this comparison, we relied solely on the templates mentioned above.

### Evaluation of usage

To further evaluate how much of these documentation approaches are used, we searched for documentation of actual datasets. We refer to them as real-world usage of the dataset documentation approaches. We followed four methods to find existing dataset documentation.

#### Method 1: References/mentions in the dataset documentation resources

We reviewed the main papers and resources associated with each documentation approach (mentioned in the “Review of dataset documentation approaches for responsible AI” section above) to check if they mentioned any datasets documented using their approach.

#### Method 2: Citation to the dataset documentation resources

We looked at resources citing the main paper(s) of each documentation approach as of March 18, 2024 with the idea that documentation created using any of these approaches may cite the associated paper.^9,11–15^ The Publish or Perish software was used to obtain a list of all such citing articles.^48^ We retained only journal articles in English accessible through the UCSD Library System. We then manually gathered the PDFs of the articles that were retained. Given the large number of these articles, we wrote a Python script in the form of a Jupyter notebook called real-world-usage.ipynb (see the **Code Availability** section) to scan the full articles for the name of the documentation approach. Specifically, we defined a set of key terms for each documentation approach, e.g. “datasheet”, “datasheets” (case insensitive) for Datasheet. If two or more key terms from such a set were found in an article citing the associated data documentation’s paper, the article was fully read to see if it contained or referred to the documentation of an actual dataset. The Python script uses packages such as PyPDF2.^49^ A list of all citing articles for each data documentation approach, including the key terms used in the code to identify articles that were fully read, is provided in the real-world-usage-method2-citations.xlsx file (see **Data Availability** section).

#### Method 3: GitHub search

We also searched for dataset documentation on GitHub, as we learned from methods 1 and 2 that many data documentation approaches are shared on GitHub. Searching for broad terms like “Datasheet” and “data card” led to a very large number of results that were mostly not relevant. To achieve better results, we selected a couple of questions or fields from each data documentation approach and searched for them within files available on GitHub. We added code to the same real-world-usage.ipynb Jupyter notebook to perform the search programmatically using the PyGitHub package.^50^ The search was performed during September 2024 with no bounds on the timeframe. We reviewed each result to check if the file returned in the search was a relevant data documentation approach or not. The search logic used for each of them, their search dates, and results are provided in the real-world-usage-method3-GitHub.xlsx file (see the **Data Availability** section).

#### Method 4: Other

We added under this method the dataset documentation we found through a non-targeted search (e.g., we were searching for Datasheets but found a Data Card).

We documented the following information in a spreadsheet called real-world-usage.xlsx (see the **Data Availability** section) for each dataset documentation we found: Method for finding (could be multiple), DOI or link of the resource mentioning the documentation, Details for finding (e.g., citation to Datasheet paper, GitHub search, etc.), Date found, Name of the dataset, DOI or link to the dataset, DOI or link to the documentation, Documentation approach (one of the five approaches of interest), Indication of whether it is for health related dataset. Here, we considered a dataset to be a health-related dataset if it included data from human subjects collected as part of a health-related study. We added code to the same real-world-usage.ipynb Jupyter notebook and used packages such as pandas^43,44^, Matplotlib^45^, seaborn^46^, and Numpy^47^ to analyze and visualize our findings, such as the number of real-world usage found for each documentation approach and how many of those were for health-related datasets

### Survey of data generators and data consumers

To understand further the usage and usefulness of these data documentation approaches in health datasets, we surveyed two categories of domain experts between July 2024 and September 2024: 1) Data generators, i.e., people who collect/generate health data, and 2) Data consumers, i.e., people who use health datasets for developing AI models. These experts were found by putting out a call for participants through various conferences and meetings, such as the 2024 Face-to-face meeting of the NIH Bridge2AI Program and the 2024 annual meeting of the Association for Research in Vision and Ophthalmology (ARVO). We also made announcements on digital platforms such as LinkedIn and Twitter. We additionally reached out to specific experts known to the authors in the AI-READI project, NIH Bridge2AI Program, and the NIH SPARC Program. We used Qualtrics software (Copyright ©2024 Qualtrics) Version June 2024 to conduct the survey (Qualtrics and all other Qualtrics product or service names are registered trademarks or trademarks of Qualtrics, Provo, UT, USA. https://www.qualtrics.com). The survey was anonymous. The survey participants were first asked to answer questions about their background and their awareness of the data documentation approaches. Then, they were asked to review a template of a randomly assigned dataset documentation approach and answer questions based on their role:

- Dataset generators were asked to rate how easy it would be to provide the information requested in each section of the dataset documentation if they were to provide it for their health dataset. They were also asked to rate the overall ease of preparing the documentation approach. A 5-point Likert scale was used for the ratings.
- Dataset consumers were asked to rate how useful the information provided in each section of the dataset documentation would be if it was provided to them when reusing a health dataset for developing AI models. They were also asked to rate the overall usefulness of the documentation approach for responsibly developing AI models. A 5-point Likert scale was used for the ratings.
- Participants who responded being both data generators and consumers were asked to review their assigned documentation approach first from a data generator’s point of view and then from a data consumer’s point of view.

After reviewing one dataset documentation approach, reviewers were asked to review another one if they wanted to, until they declined to continue or reviewed all five. A copy of the survey and raw survey responses are in the survey.pdf and survey-results.xlsx files, respectively, included in the dataset associated with this work (see **Data Availability** section). We wrote a Python script in the form of a Jupyter notebook called survey-analysis.ipynb (see the **Code Availability** section) to analyze the survey results exported from Qualtrics and generate relevant plots and tables using Python packages such as pandas^43,44^, Matplotlib^45^, seaborn^46^, and Numpy^47^.

## Declarations

### Human Ethics and Consent to Participate declarations

not applicable.

### Ethics Approval declaration

This study was reviewed by the University of California San Diego Institutional Review Board (IRB) and was granted exemption from IRB review.

### Author Contributions

A.H.- wrote the main manuscript text, collected the data, analyzed the data, reviewed the manuscript, L. H.-wrote the main manuscript text, collected the data, analyzed the data, reviewed the manuscript, K.U.S.-wrote the main manuscript text, collected the data, developed the code, analyzed the data, reviewed the manuscript F.G.P.K-wrote the main manuscript text, collected the data, reviewed the manuscript A.K.-wrote the main manuscript text, collected the data, reviewed the manuscript K.S.-collected the data, reviewed the manuscript, S.S.- collected the data, reviewed the manuscript, C. N.-contributed to the methodology, reviewed the manuscript, S.L.B.-contributed to the methodology, reviewed the manuscript C.S. L-contributed to the methodology, reviewed the manuscript, A.Y.L-contributed to the methodology, reviewed the manuscript, B.P.- designed the study, wrote the main manuscript text, collected the data, developed the code, analyzed the data, reviewed the manuscript.

## References

1. Muehlematter, U. J., Daniore, P. & Vokinger, K. N. Approval of artificial intelligence and machine learning-based medical devices in the USA and Europe (2015-20): a comparative analysis. Lancet Digit. Health 3, e195–e203 (2021).

2. Sidey-Gibbons, J. A. M. & Sidey-Gibbons, C. J. Machine learning in medicine: a practical introduction. BMC Med. Res. Methodol. 19, 64 (2019).

3. Chen, I. Y. et al. Ethical machine learning in healthcare. Annu. Rev. Biomed. Data Sci. 4, 123–144 (2021).

4. Ibrahim, H., Liu, X., Zariffa, N., Morris, A. D. & Denniston, A. K. Health data poverty: an assailable barrier to equitable digital health care. Lancet Digit. Health 3, e260–e265 (2021).

5. Schwartz, R., et al. Towards a Standard for Identifying and Managing Bias in Artificial Intelligence. https://doi.org/10.6028/NIST.SP.1270 (2022) doi:10.6028/nist.sp.1270.

6. Seyyed-Kalantari, L., Zhang, H., McDermott, M. B. A., Chen, I. Y. & Ghassemi, M. Underdiagnosis bias of artificial intelligence algorithms applied to chest radiographs in under-served patient populations. Nat. Med. 27, 2176–2182 (2021).

7. Daneshjou, R. et al. Disparities in dermatology AI performance on a diverse, curated clinical image set. Sci Adv 8, eabq6147 (2022).

8. Arora, A. et al. The value of standards for health datasets in artificial intelligence-based applications. Nat. Med. 29, 2929–2938 (2023).

9. Gebru, T. et al. Datasheets for datasets. Commun. ACM 64, 86–92 (2021).

10. Alderman, J. E. et al. Tackling algorithmic bias and promoting transparency in health datasets: the STANDING Together consensus recommendations. Lancet Digit. Health 7, e64–e88 (2025).

11. Holland, S., Hosny, A., Newman, S., Joseph, J. & Chmielinski, K. The Dataset Nutrition Label: A Framework To Drive Higher Data Quality Standards. arXiv [cs.DB] (2018).

12. Chmielinski, K. S., et al. The Dataset Nutrition Label (2nd Gen): Leveraging Context to Mitigate Harms in Artificial Intelligence. arXiv [cs.LG] (2022).

13. Hutchinson, B. et al. Towards Accountability for Machine Learning Datasets: Practices from Software Engineering and Infrastructure. in Proceedings of the 2021 ACM Conference on Fairness, Accountability, and Transparency 560–575 (Association for Computing Machinery, New York, NY, USA, 2021).

14. Rostamzadeh, N. et al. Healthsheet: Development of a Transparency Artifact for Health Datasets. in Proceedings of the 2022 ACM Conference on Fairness, Accountability, and Transparency 1943–1961 (Association for Computing Machinery, New York, NY, USA, 2022).

15. Pushkarna, M., Zaldivar, A. & Kjartansson, O. Data Cards: Purposeful and Transparent Dataset Documentation for Responsible AI. in Proceedings of the 2022 ACM Conference on Fairness, Accountability, and Transparency 1776–1826 (Association for Computing Machinery, New York, NY, USA, 2022).

16. Ganapathi, S. et al. Tackling bias in AI health datasets through the STANDING Together initiative. Nat. Med. 28, 2232–2233 (2022).

17. AI-READI Consortium. AI-READI: rethinking AI data collection, preparation and sharing in diabetes research and beyond. Nat. Metab. 6, 2210–2212 (2024).

18. The Data Nutrition Project. https://datanutrition.org.

19. The Data Cards Playbook. https://sites.research.google/datacardsplaybook.

20. Gebru, T., et al. Datasheets for Datasets Preprint. arXiv [cs.DB] (2018) doi:10.48550/arXiv.1803.09010.

21. DNP Label Maker. https://labelmaker.datanutrition.org.

22. Hutchinson, B., et al. Towards Accountability for Machine Learning Datasets: Practices from Software Engineering and Infrastructure. arXiv [cs.LG] (2020).

23. Healthsheet: Development of a Transparency Artifact for Health Datasets. https://dl.acm.org/doi/fullHtml/10.1145/3531146.3533239.

24. Pushkarna, M., Zaldivar, A. & Kjartansson, O. Data Cards: Purposeful and Transparent Dataset Documentation for Responsible AI. arXiv [cs.HC] (2022).

25. Healthsheet: Development of a Transparency Artifact for Health Datasets. https://dl.acm.org/doi/fullHtml/10.1145/3531146.3533239.

26. templates/DataCardsExtendedTemplate.pdf at Main · PAIR-Code/datacardsplaybook. (Github).

27. Bandrowski, A. et al. SPARC Data Structure: Rationale and Design of a FAIR Standard for Biomedical Research Data. bioRxiv 2021.02.10.430563 (2021) doi:10.1101/2021.02.10.430563.

28. Marroquin, C. et al. SODA: Software to support the curation and sharing of FAIR autonomic nervous system data. J. Open Source Softw. 9, 6140 (2024).

29. Marroquin, A., et al. SODA (Software to Organize Data Automatically) for SPARC. (Zenodo, 2024). doi:10.5281/ZENODO.5565455.

30. Heinke, A., et al. Dataset: Dataset Documentation for AI Paper. Zenodo 10.5281/ZENODO.17308743 (2025).

31. Patel, B., Soundarajan, S., Ménager, H. & Hu, Z. Making Biomedical Research Software FAIR: Actionable step-by-step guidelines with a user-support tool. Sci. Data 10, 557 (2023).

32. Portillo, D., Soundarajan, S. & Patel, B. Codefair. (Zenodo, 2024). doi:10.5281/ZENODO.13376616.

33. Simpkins, K. & Patel, B. Code: Dataset Documentation for AI Paper. (Zenodo, 2025). doi:10.5281/ZENODO.17308727.

34. Yang, X., Liang, W. & Zou, J. Navigating dataset documentations in AI: A large-scale analysis of dataset cards on Hugging Face. arXiv [cs.LG] (2024) doi:10.48550/ARXIV.2401.13822.

35. Srinivasan, R., et al. Artsheets for Art Datasets. (2021).

36. Jain, N., et al. A standardized machine-readable dataset documentation format for responsible AI. arXiv [cs.IR] (2024) doi:10.48550/ARXIV.2407.16883.

37. Roman, A. C., et al. Open datasheets: Machine-readable documentation for open datasets and responsible AI assessments. arXiv [cs.LG] (2023) doi:10.48550/ARXIV.2312.06153.

38. Siddik, M. & Pandit, H. J. Datasheets for healthcare AI: A framework for transparency and bias mitigation. arXiv [cs.CY] (2025) doi:10.48550/ARXIV.2501.05617.

39. Wang, W., et al. MiniLM: Deep self-attention distillation for task-agnostic compression of pre-trained transformers. arXiv [cs.CL] (2020) doi:10.48550/ARXIV.2002.10957.

40. sentence-transformers/all-MiniLM-L6-v2 · Hugging Face. https://huggingface.co/sentence-transformers/all-MiniLM-L6-v2/.

41. Reimers, N. & Gurevych, I. Sentence-BERT: Sentence embeddings using Siamese BERT-networks. arXiv [cs.CL] (2019) doi:10.48550/ARXIV.1908.10084.

42. Reimers, N. & Gurevych, I. Making monolingual sentence embeddings multilingual using knowledge distillation. arXiv [cs.CL] (2020) doi:10.48550/ARXIV.2004.09813.

43. The pandas development team. Pandas-Dev/pandas: Pandas. (Zenodo, 2024). doi:10.5281/ZENODO.3509134.

44. McKinney, W. Data Structures for Statistical Computing in Python. in Proceedings of the Python in Science Conference 56–61 (SciPy, 2010).

45. Hunter, J. D. Matplotlib: A 2D Graphics Environment. Comput. Sci. Eng. 9, 90–95 (2007).

46. Waskom, M. seaborn: statistical data visualization. J. Open Source Softw. 6, 3021 (2021).

47. Harris, C. R. et al. Array programming with NumPy. Nature 585, 357–362 (2020).

48. Harzing, A. W. Perish or Publish. https://harzing.com/resources/publish-or-perish (2007).

49. Fenniak, M., et al. The PyPDF2 library. Preprint at https://pypi.org/project/PyPDF2/ (2022).

50. PyGithub: Typed Interactions with the GitHub API v3. (Github).

51. Datasheet-for-Dataset-Template: Template for Datasheet for Datasets. (Github).

52. Datasheet for dataset template. https://www.overleaf.com/latex/templates/datasheet-for-dataset-template/jgqyyzyprxth.

53. templates/Healthsheet Template.docx at Main · PAIR-Code/datacardsplaybook. (Github).

54. The Data Cards Playbook. https://sites.research.google/datacardsplaybook.

55. Datacardsplaybook: The Data Cards Playbook Helps Dataset Producers and Publishers Adopt a People-Centered Approach to Transparency in Dataset Documentation. (Github).

